# A glia cell dependent mechanism at a peripheral nerve plexus critical for target-selective axon regeneration

**DOI:** 10.1101/2023.01.05.522786

**Authors:** Lauren J Walker, Camilo Guevara, Koichi Kawakami, Michael Granato

## Abstract

A critical step for functional recovery from peripheral nerve injury is for regenerating axons to connect with their pre-injury targets. Reestablishing pre-injury target specificity is particularly challenging for limb-innervating axons as they encounter a plexus, a network where peripheral nerves converge, axons from different nerves intermingle, and then re-sort into target-specific bundles. Here, we examine this process at a plexus located at the base of the zebrafish pectoral fin, equivalent to tetrapod forelimbs. Using live cell imaging and sparse axon labeling, we find that regenerating motor axons from three nerves coalesce into the plexus. There, they intermingle and sort into distinct branches, and then navigate to their original muscle domains with high fidelity that restores functionality. We demonstrate that this regeneration process includes selective retraction of mistargeted axons, suggesting active correction mechanisms. Moreover, we find that Schwann cells are enriched and associate with axons at the plexus, and that Schwann cell ablation during regeneration causes profound axonal mistargeting. Our data provide the first real time account of regenerating vertebrate motor axons navigating a nerve plexus and reveal a previously unappreciated role for Schwann cells to promote axon sorting at a plexus during regeneration.

## INTRODUCTION

Axons of the peripheral nervous system (PNS) have significant capacity to regenerate after injury from chemical or mechanical insults. To achieve functional recovery, regenerating motor axons face the challenge of reconnecting with their original muscle targets. In mammals, motor axon targeting errors during regeneration are common and can cause long term deficits (reviewed in:^1,2^). Recently discovered pro-regenerative molecular pathways enhance axon growth during regeneration^3–7^, however these manipulations frequently lack the spatial cues that direct axons to their original targets and therefore limit functional recovery. Furthermore, while we now have a broad understanding of the developmental cues that shape the nervous system, there is growing evidence that axon regeneration is not simply a recapitulation of development, but that it requires unique injury-dependent signals^8–10^.

Target specificity is particularly challenging when regenerating axons navigate through a series of choice points. The brachial plexus, the complex network of peripheral nerves at the base of the tetrapod forelimb, is one such region. Limb-innervating peripheral nerve axons exit from the spinal cord within discrete nerves which converge at the brachial plexus. There, axons from several nerves intermingle and then sort into target-specific bundles prior to innervating certain muscles in the forelimb. Recent work has revealed several molecular mechanisms that instruct axon guidance decisions at choice points during regeneration^8,9,11,12^, yet the molecular and cellular mechanisms that enable proper axon navigation through a plexus followed by multiple choice points to innervate the proper muscle targets are largely unknown. Denervated Schwann cells, which upregulate trophic factors and form tracks, called Bands of Bungner, upon which regenerating axons grow^13^, are one cell type that could function in this process.

The larval zebrafish pectoral fin, evolutionarily analogous to tetrapod forelimbs^14^, has stereotyped motor innervation. At 5 days post fertilization (dpf), pectoral fins are comprised of two antagonistic muscles, the abductor and adductor. The fin musculature is innervated by 4 distinct motor nerves, which we refer to here as nerves 1-4, with cell bodies in anterior spinal cord segments 3 through 6 **(Fig 1A)**^15,16^. Motor nerves grow through axial trunk muscle and converge at the dorsal plexus located at the dorsal anterior edge of the fin (nerves 1-3) or the ventral plexus located at the ventral anterior edge of the fin (nerve 4). At these plexuses, axons sort between the abductor or adductor muscles **(Fig 1B)**^16^ and then segregate into target-selective bundles (**Fig 1 A,C)**. Motor axons innervate the fin musculature topographically, such that neurons from the more anterior spinal cord segment 3 (nerve 1) innervate the dorsal region of fin musculature, while neurons from spinal segments 4 and 5 (nerves 2 and 3) innervate the middle region, and neurons from spinal segment 6 (nerve 4) innervate the ventral region of the musculature (**Fig 1A)**^16,17^. Consequently, to connect to their correct muscle fibers, fin motor axons navigate a series of stepwise choice points. They first sort between muscles at the plexus and then, at subsequent choice points within the fin musculature, axons select a path to innervate the proper muscle domain. Thus, the complex pectoral fin motor innervation pattern combined with the genetic-tractability, optical transparency suitable for live imaging, and the ability to measure fin movement as a functional readout of regeneration, make the larval zebrafish pectoral fin an ideal vertebrate system to interrogate mechanisms of target-selective axon regeneration.

**Figure 1:**
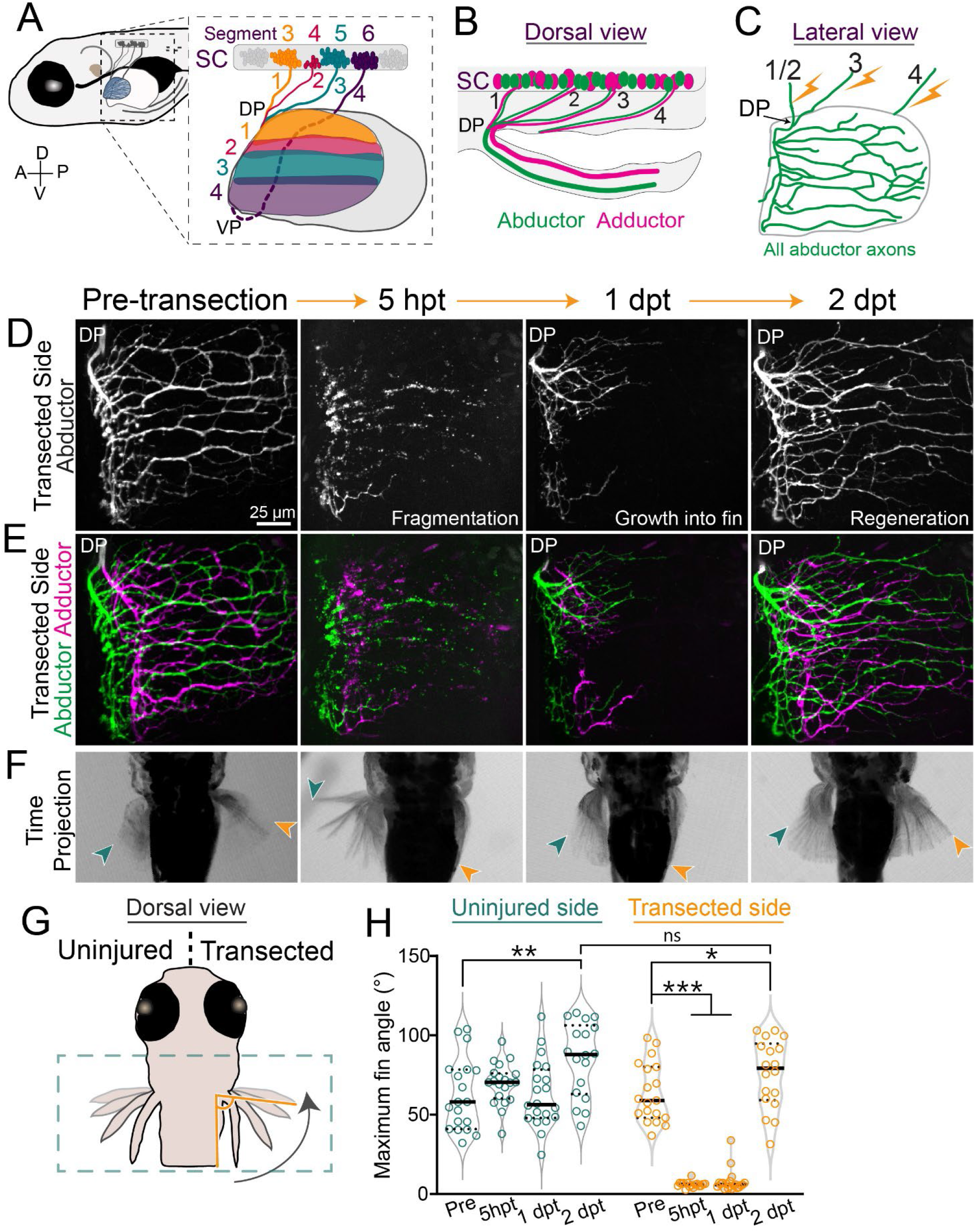
Pectoral fin motor axons regenerate robustly and reform functional synapses. A) Schematic of a 5 day post fertilization (dpf) larval zebrafish. Inset shows motor pools from spinal cord (SC) segments 3-6 that form pectoral fin nerves 1-4 in the body wall. Axons sort at a dorsal (DP) or ventral plexus (VP) and innervate the musculature of the pectoral fin topographically. Innervation domains are labeled 1-4 and shown in corresponding colors. B) Dorsal view. Motor neurons in spinal cord segments innervate either the abductor or adductor muscle. C) Lateral view. Schematic of abductor innervation of the pectoral fin. Nerves were transected using a laser in the locations shown with the lightning bolts. D-E) Images of maximum projections of fin motor innervation labeled with *tubb:dsRed* from the transected side. This example is from the same animal through regeneration. At 5 hours post transection (hpt), axons have fragmented. At 1 day post transection (dpt) axons have started to grow into the fin. At 2 dpt, axons have regenerated. D) Abductor muscle innervation. E) Abductor and adductor innervation pseudo-colored in green and magenta, respectively. F) Corresponding time projections of spontaneous pectoral fin movements. The region shown is indicated by the green dotted box in G. Only the nerves of the right fin were injured. At 5 hpt, the injured fin does not move. At 1 dpt, axons have just begun to grow into the fin but the injured fin still does not move. However, by 2 dpt axons have regenerated and the injured fin can move again. The green and orange arrows point to the maximum fin position for the uninjured and transected sides, respectively. G) The maximum angle of the tip of the fin compared to the body wall was measured during spontaneous fin movements. The box indicates the anatomical region imaged in F. H) Quantification of the maximum angle of the uninjured and transected fins pre-transection through regeneration. Each dot represents one fish and one movement. One-way ANOVA. *p=0.04, **p<0.005, ***p<0.0001, ns = not significant.

Here, we use larval zebrafish to visualize the multistep process of axonal regeneration through a peripheral nerve plexus in real time, and to determine the role of local glia cells in this process. We demonstrate that within two days of complete motor nerve transection, motor axons regenerate robustly to reestablish functional synapses. We find that individual axons faithfully sort at the plexus to reinnervate their original muscle fiber targets and that mistargeted axons are selectively retracted to correct targeting errors. Finally, we demonstrate that Schwann cells are required for regenerating axons to properly navigate through the plexus, in part by preventing axonal mistargeting to the incorrect muscle. Combined, this work reveals a previously unappreciated role for Schwann cells to ensure appropriate axon sorting at a plexus, thereby promoting target-selective axon regeneration.

## RESULTS

### Pectoral fin motor axons regenerate robustly

To identify the mechanisms that guide regenerating peripheral nerve axons through a plexus, we first observed axons during the process of regeneration. To visualize motor nerves that innervate the pectoral fin we used transgenic *Xla.Tubb:dsRed* (hereafter referred to as *Tubb:dsRed*) larvae. At 5 dpf, pectoral fin targeting motor axons have established an elaborate innervation field across the abductor and adductor musculature of the fin (**Fig 1D-E**). We used a laser to transect nerves 1/2 and 3 dorsal to the dorsal plexus and nerve 4 at the same approximate area within the body wall **(Fig 1C)**. This laser transection strategy yielded complete motor denervation of the pectoral fin while leaving the fin itself uninjured. Within 2-3 hours post transection (hpt), the distal portion of transected axons began to bleb and axons fragmented by 5 hpt. Following an initial period of stasis, axons initiated growth and navigated the dorsal plexus at 12 hpt ± 2.6 hours (n=9) to sort between the abductor and adductor muscles. Axons exit the dorsal or ventral plexus as a single fascicle prior to segregating into discrete bundles. In this study, we focused predominately on the dorsal plexus and hereafter all results pertain to axon regeneration through the dorsal plexus. By 24 hpt, these discrete bundles were apparent as regenerating axons extended partially across the musculature. Axon regeneration proceeded rapidly, with axon regrowth largely completed by 48 hpt (2 days post transection (dpt)) **(Fig 1D-E)**. Thus, pectoral fin motor axons regenerate robustly to reestablish the complex innervation patterning.

### Pectoral fin motor axons achieve functional regeneration

To determine the degree of functional regeneration, we measured fin movements prior to and following nerve transection. Larval zebrafish pectoral fins perform spontaneous movements of alternating pectoral fins that are dependent on motor axon innervation^18^. To determine if and to what extent pectoral fin innervating motor axons achieve functional recovery, we used high-speed imaging at millisecond resolution to record spontaneous pectoral fin movements. From these movies, we then determined the maximum fin movement angle **(Fig 1F-H)**. Concurrently, we also imaged fin motor axon innervation in the same animal throughout the regeneration process and then correlated fin movement with the extent of innervation. Transecting nerves that innervate the right pectoral fin while sparing those that innervate the left pectoral fin provided an internal control. Prior to nerve transection, left and right pectoral fins moved rhythmically (mean maximum angles for left fin 62.5 ± 22.7 degrees and for right fin 63.7 ± 18.73 degrees, n=19; p= 0.856, unpaired t-test) **(Fig 1H, Mov 1**). Five hours after nerve transection, motor axons that innervated the right pectoral fin had fragmented and all movements of the right, transected fin ceased, whereas the uninjured left fin continued to move. At 1 dpt, regenerating axons had partially grown back into the dorsal region of the right fin, yet the fin still failed to move. In contrast, at 2 dpt axons had fully regrown across the fin musculature and the injured fin performed spontaneous movements comparable to the uninjured fin (mean maximum angle at 2 dpt for uninjured fin 85.0 ± 23.3 degrees and for transected fin 75.8 ± 21.8 degrees; p= 0.233, unpaired t-test) (**Fig 1G-H)**. This recovery of movement demonstrates that pectoral fin motor axons do not only regrow robustly towards their original targets, but that they also reform functional synapses.

### Single axon labeling reveals target specificity

We next examined the specificity with which regenerating axons grew back to their original muscle domains that were established during development. For this, we sparsely labeled axons using *mnx1:mKate* in the context of the entire population of labeled motor neurons (*mnx1:GFP*) (**Fig 2A)**. This approach allows for a direct comparison of the trajectory of one or only a few axons before transection and after regeneration. Two days after complete transection of all pectoral fin innervating motor nerves, sparsely labeled axons had correctly navigated many choice points to reestablish an innervation pattern similar to their pre-injury route (**Fig 2 B-C)**. Next, we examined the frequency by which regenerating axons reinnervated their original muscle (abductor vs adductor) and their original topographic domains. We found that 94.5% of sparsely labeled axons re-occupied their original muscle (n=52/55 to the correct muscle) (**Fig 2D**). To determine the specificity of axon targeting within the musculature we divided the fin into four domains based on axon targeting patterns (**Fig 2E**). We found that 85.5% of axons reinnervate their original domains (n=47/54 to the correct domain) (**Fig 2F**). Thus, following complete nerve transection, pectoral fin motor axons reinnervate their original muscle and muscle domains with high specificity. This specificity strongly suggests the existence of robust mechanisms during regeneration that mediate axon sorting at the plexus between the abductor versus the adductor muscle, and precise domain targeting of axons throughout the fin musculature.

**Figure 2:**
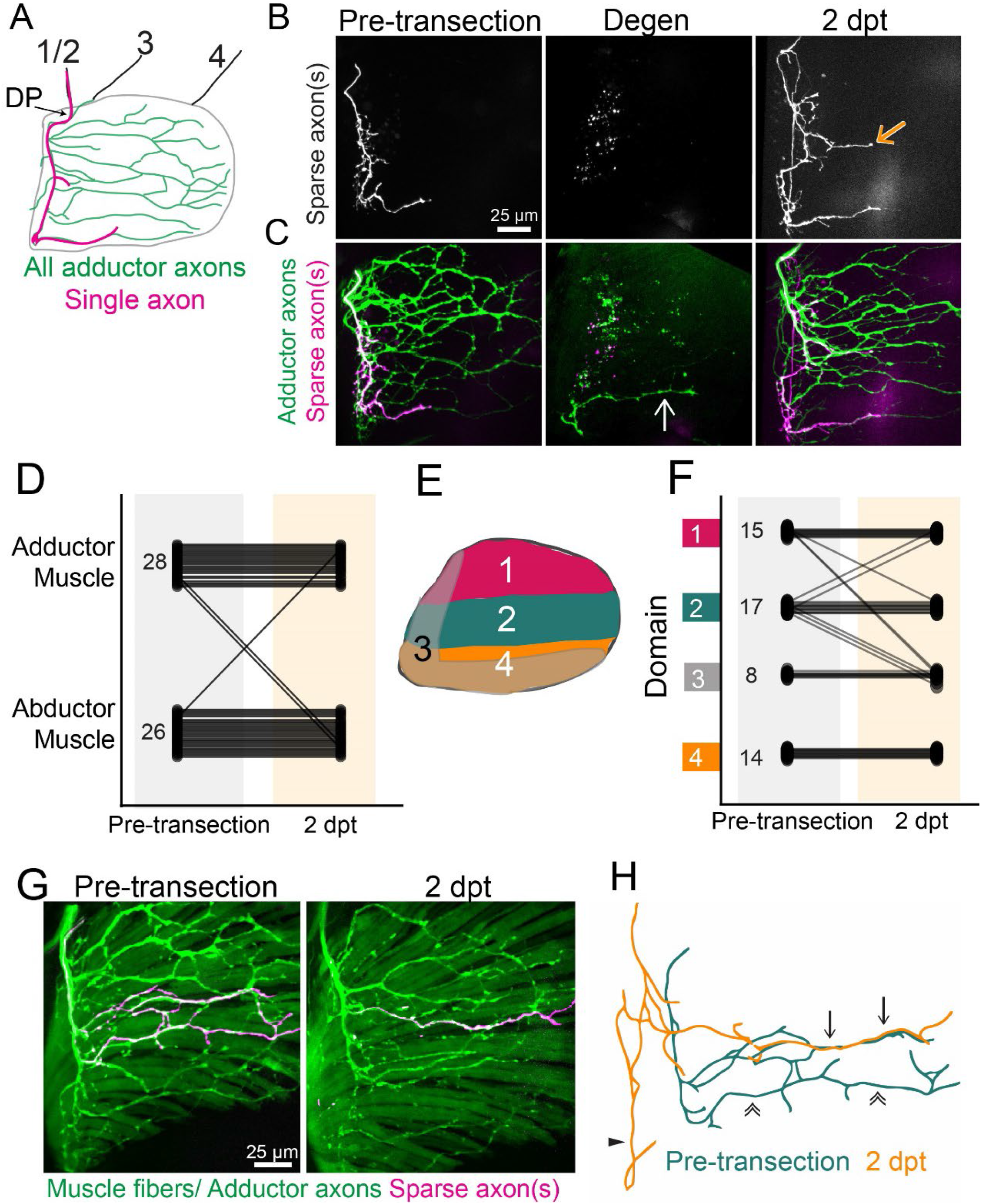
Target-selective regeneration of motor axons. A) Schematic of a lateral view of pectoral fin abductor muscle motor innervation. An example trajectory of a single axon is shown in magenta. DP labels dorsal plexus, motor nerves in the body wall are labeled 1-4. B-C) A timecourse of sparsely labeled axon(s) showing the innervation pattern pre-transection, after axon degeneration, and at 2 days post transection (dpt). Sparsely labeled axons are labeled in white (B) and magenta (C) and all adductor motor axons are labeled in green (C). The white arrow points to a fascicle that did not degenerate, and the orange arrow points to ectopic growth during regeneration. D) Quantification of sparsely labeled axon muscle localization pre-transection and after regeneration. n= 28 abductor and 26 adductor pre-injury. E) Schematic defining domains for axon domain scoring. For category 3, axons entered the fin at the dorsal plexus but innervated the ventral region of the fin, overlapping with domain 4 (like the single axon in A). F) Quantification of fin domain localization. Small numbers on pre-transection data represent n’s. G) Example of sparsely labeled axons (magenta) that form new trajectories and re-establish previous trajectories during regeneration with both motor axons (*mnx1:GFP*) and muscle fibers (*α-actin:GFP*) labeled. H) The original (green) and regenerated (orange) trajectories of the sparsely labeled axons in G. Here, part of the pre-injury and regenerated trajectory can be overlayed precisely (arrows). Additionally, an axon does not follow its original trajectory (double arrows) but instead is mistargeted along the base of the fin (triangle). See also Supp. Fig. 1.

Both the abductor and adductor muscles are comprised of ~50 muscle fibers arranged longitudinally across the fin^19^. To determine how precise fin motor axon targeting is during regeneration, we next asked if labeled motor axons reinnervate their original muscle fibers. For this, we used larvae in which all motor neurons and muscle fibers were labeled with green fluorescent protein (GFP) (*mnx1:GFP; α-actin:GFP*), and then used *mnx1:mKate* to sparsely label axons. The position and morphology of muscle fibers remained consistent through the course of the experiment, allowing them to serve as landmarks. Using this framework, after injury, sparsely labeled motor axons regenerated to their original muscle fiber targets with high but not perfect precision (**Fig 2 G, Supp Fig 1A-B**). Specifically, we observed that during the regeneration process some terminal axonal branches vacated their original, pre-injury muscle fibers and that some now occupied new muscle fibers. The example shown in figure **2 G-H** illustrates the variability of axon targeting we observe during regeneration. Here, a section of the route from sparsely labeled regenerated axons was distinct from the pattern prior to injury indicating that during regeneration axons sometimes form divergent projections. However, the distal segment of the regenerated axon pattern can be directly overlayed on the pre-injury axon path, demonstrating that the distal end of this axon followed the same route it had prior to injury (of 9 muscles with sparse axon labeling: 3 had labeled axons that terminated on their original muscle fibers, 3 had axons that partially grew across their original muscle fibers but terminated elsewhere, 2 had labeled axons that failed to regrow, and 1 had labeled axons that innervated the original domain but not the original muscle fibers). Thus, pectoral fin motor axon regeneration is remarkably precise, such that axons preferentially re-innervate their original fin domains, with specificity as accurate as their original muscle fibers.

### Correction of mistargeted axons during regeneration

The small but consistent fraction of regenerating axons that displayed mistargeting prompted us to examine if mistargeted axons remained stable or were corrected. For this, we employed transgenic *zCrest2-hsp70:GFP* (referred to hereafter as *zCrest:GFP*) larvae in which motor axons that project to the abductor muscle are selectively labeled^20^. Prior to nerve injury, zCrest:GFP-labeled axons almost exclusively innervated the abductor muscle (n=38/41, 92.7% pectoral fins), and only a small fraction of fins (n=3/41, 7.3% pectoral fins) contained any zCrest:GFP-labeled axons on the adductor muscle. In contrast, at 2 dpt, zCrest:GFP-labeled axons were present on the adductor muscle in 97.37% of pectoral fins examined (n= 40/41). Despite the presence of these mistargeted axons, the proportion of correctly targeted axons on the abductor muscle was much higher than mistargeted axons on the adductor muscle, suggesting that at the plexus only a small population of axons fail to select their original muscle. This is consistent with our sparse labeling approach, which estimates that ~4% of axons mistargeted to the incorrect muscle (**Fig 2C)**.

To determine if mistargeted axons are subsequently corrected, we assessed static timepoints during the regeneration process. In *zCrest:GFP; Tubb:dsRed* larvae, all axons are labeled with dsRed fluorescent protein and abductor-specific motor axons are labeled with GFP. Prior to axon injury, zCrest:GFP-labeled axons project exclusively to the abductor muscle (**Fig 3A-C)**. Following transection of all fin motor nerves, the disconnected distal portion of motor axons on both the abductor and the adductor muscle fragmented, and axonal debris was partially cleared at 7 hpt (**Fig 3D)**. At 20 hpt, zCrest:GFP-labeled axons were present on both the abductor and adductor muscles, suggesting that in the early phases of regeneration, axons are competent to grow on both muscles independent of their pre-injury targets (**Fig 3E)**. The extent of mistargeted axon growth on the adductor muscle varied between fish (804 ± 508 microns, minimum 134 microns and maximum 1497 microns of mistargeted axon growth, n=9 fins). At 30 hpt some mistargeted zCrest:GFP-labeled fascicles began to bleb or had already retracted, whereas other mistargeted fascicles continued to extend into the adductor musculature (**Fig 3F)**. By 50 hpt, more of the mistargeted zCrest:GFP-labeled fascicles had been retracted; however, there were also misprojected fasicles that stabilized and persisted (**Fig 3G)**. Retraction events of mistargeted zCrest:GFP-labeled axons occurred in 100% of animals examined (15/15 pectoral fins). At the same time that misprojected axons retracted on the adductor muscle, the correctly targeted zCrest:GFP-labeled axons on the abductor muscle increased in complexity (**Fig 3 E-G**). Thus, early in regeneration, axon mistargeting at the plexus is common, yet over time misprojected axons are corrected.

**Figure 3:**
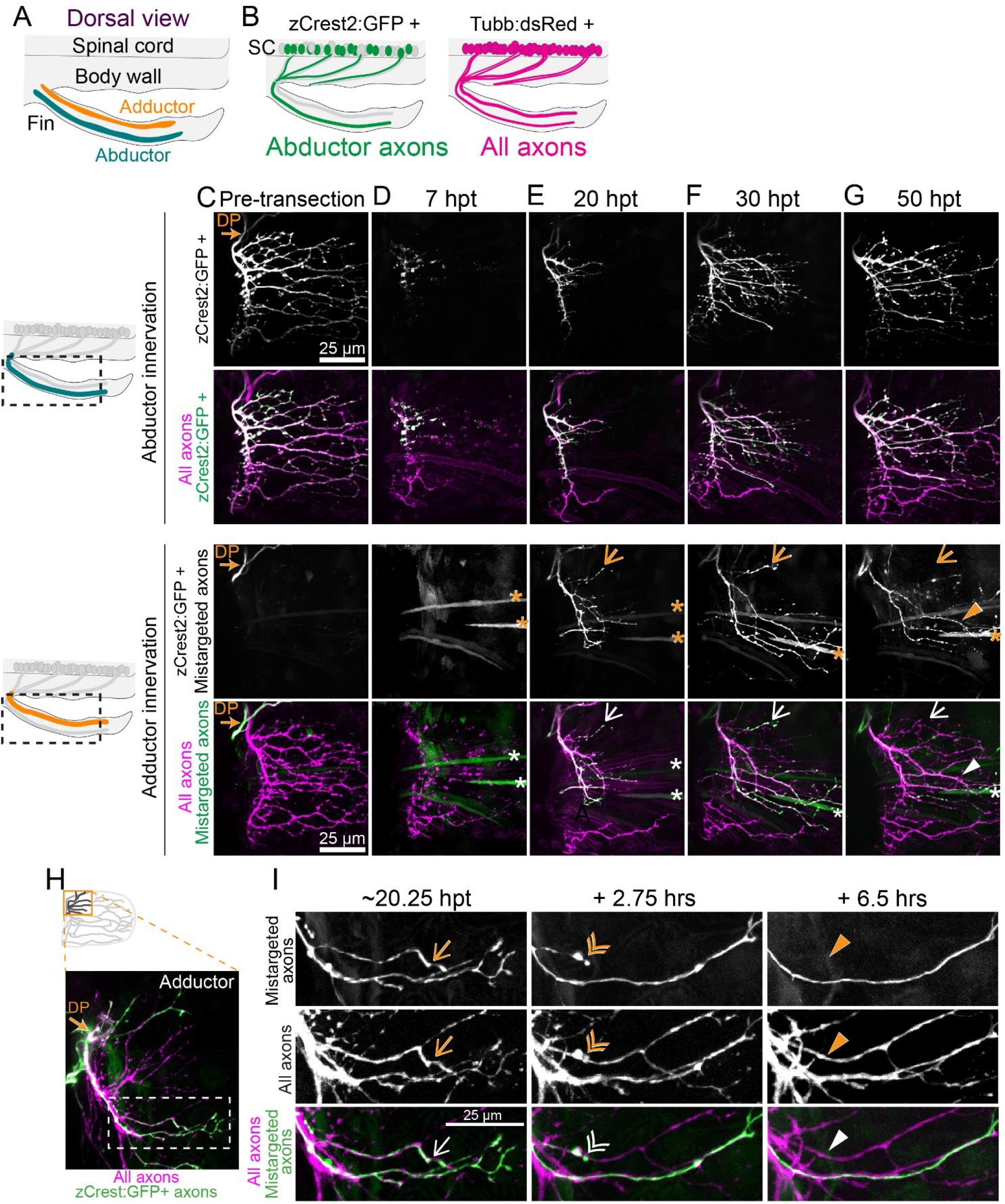
Mistargeted axons are selectively retracted. A) Dorsal view schematic labeling abductor and adductor musculature of the fin. B) Schematic of zCrest:GFP+ motor neurons, which project to the abductor muscle and *Tubb:dsRed*+ motor neurons, which project to both abductor and adductor muscles. zCrest:GFP+ motor neurons are also labeled with *Tubb:dsRed*. C-G) Timecourse of regeneration of innervation on abductor (top) and adductor (bottom) musculature. Schematics on the left show the area included in maximum projections. Arrows point to mistargeted axons that will retract. Asterisks indicate muscle fibers also labeled by *zCrest:GFP*. C) zCrest:GFP+ axons are not present on the adductor muscle before axon transection. DP labels dorsal plexus. D) At 7 hpt, axons have fragmented. E) At 20 hpt, regenerating *Zcrest2:GFP*+ axons are present on both the abductor and adductor muscle. F-G) At 30 and 50 hpt some Zcrest2:GFP*+* mistargeted axons have retracted (arrow) whereas other mistargeted axons persist (triangle). H) Maximum projection of axon regeneration onto adductor muscle at 20.25 hpt. The boxed region is expanded in I. zCrest:GFP+ axons are mistargeted onto adductor muscle. I) Images from timelapse imaging of axon regeneration onto adductor muscle. *zCrest:GFP* labels mistargeted axons whereas axons that are only magenta (labeled with *Tubb:dsRed*) are correctly targeted to adductor muscle. Example of a mistargeted axon that is present at 20.25 hpt (arrow), retracting at +2.75 hours (double arrowhead), and gone by +6.5 hours (filled triangle).

Finally, we examined the dynamic nature of axon mistargeting and retractions on the adductor muscle. By using *zCrest:GFP; Tubb:dsRed* double transgenic larvae, we can distinguish between zCrest:GFP-negative; dsRed-positive axons that correctly target to the adductor muscle and zCrest:GFP-positive; dsRed-positive axons that are presumptive mistargeted axons on the adductor muscle. Using live imaging, we observed the process by which mistargeted axons are selectively retracted. On the adductor muscle, zCrest:GFP-negative, dsRed-positive axons grew robustly and branched to tile on their appropriate target muscle. Concomitantly, after mis-sorting at the dorsal plexus onto the adductor muscle, mistargeted zCrest:GFP-labeled axons also grew robustly in the early phase of regeneration. These mistargeted axons grew within all fin domains and formed elaborate branches, similar to dsRed-positive axons. However, after an initial phase of robust growth, some mistargeted zCrest:GFP-labeled axons stopped extending, formed a retraction bulb, and retracted. The timing of these retraction events, which occurred at approximately 19 ± 2.6 hpt (n = 25 retraction events, timelapse duration <30 hpt), was consistent across pectoral fins. This retraction was specific to mistargeted axons because correctly targeted dsRed-positive adductor axons within the same fascicle remained stable and continued to grow (**Fig 3 H-I, Mov 2)**. Consistent with the static timecourse of retractions **(Fig 3 A-G)**, retraction events occurred in all fins (n = 9/9), but not all mistargeted axons within these fins were corrected. Thus, our data reveal the existence of mechanisms active during regeneration that selectively correct mistargeted axons.

### Schwann cells in the pectoral fin associate with regenerating axons

While Schwann cells associate closely with motor axons in the larval zebrafish trunk^21^, whether they are present in the pectoral fin has not been reported. To answer this question, we used the *37A:Gal4, UAS:EGFP* line^22^ to drive GFP expression in Schwann cells. Within the pectoral fin, *37A:Gal4, UAS:EGFP* labels cells with a rounded cell body and long, narrow processes. By co-labeling the fluorescent marker EGFP with the nuclear marker DAPI, we counted 12 ± 2.1 (n=8 fins) Schwann cells in the pectoral fin. Over twice as many Schwann cells reside within the abductor muscle than the adductor muscle at 5-7 dpt (abductor: mean 7.3 ± 2.4 Schwann cells; adductor: mean 2.6 ± 1.5 Schwann cells; n=8 fins) (**Fig 4A-B)**. Within the fin, most motor axons are closely associated with these Schwann cells **(Fig 4C)**. Schwann cells also reside in the vicinity of fin motor nerves located in the body wall (**Fig 4D).** At the dorsal plexus, Schwann cells completely enwrapped motor axon bundles **(Fig 4E-F)** (1.8 ± 1 Schwann cells at the dorsal plexus; n=6). Thus, prior to injury, Schwann cells are present throughout the pectoral fin and associate closely with motor axons.

**Figure 4:**
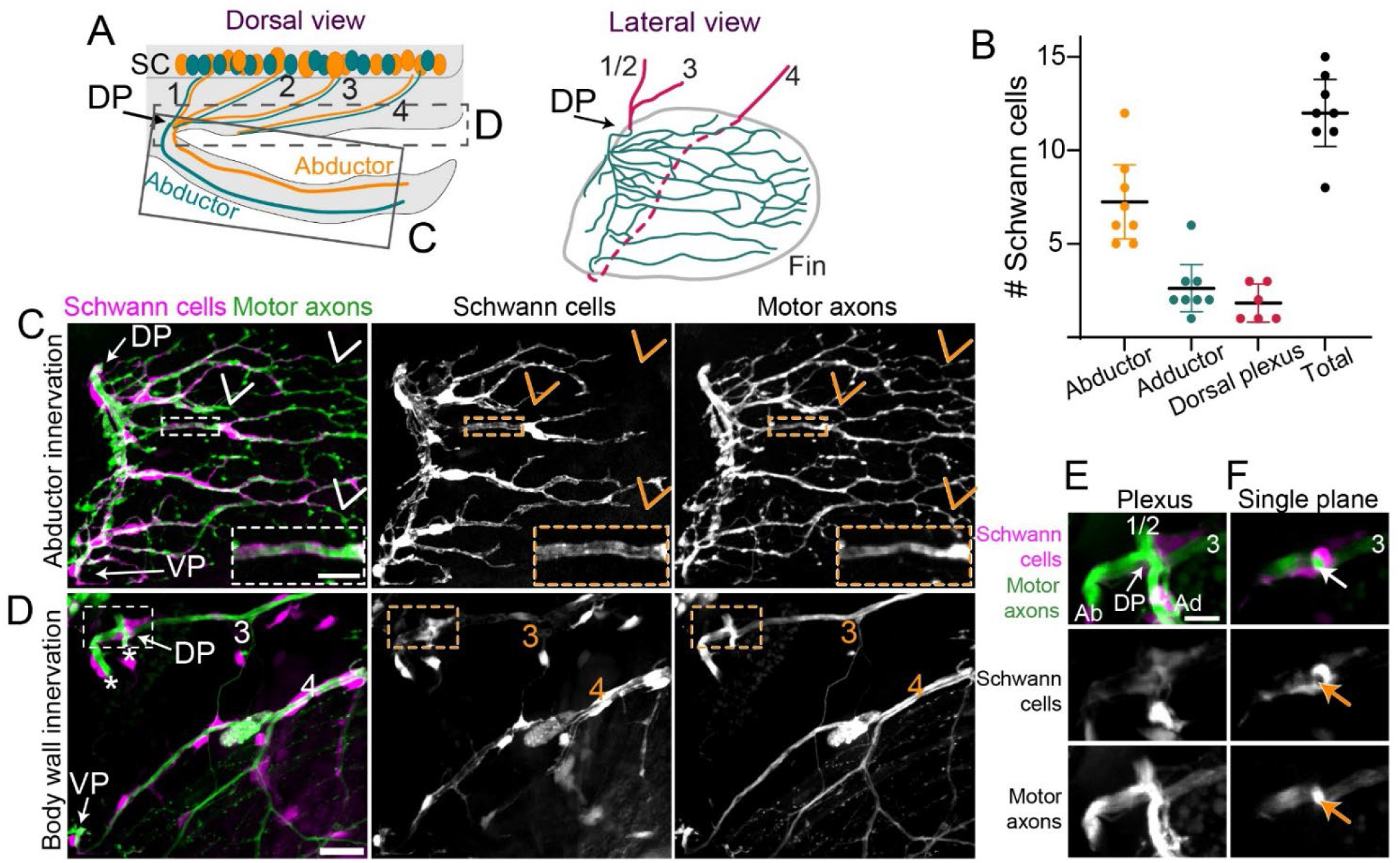
Schwann cells tightly associate with axons in the pectoral fin. A) Schematic of a dorsal view of a larval zebrafish. In the spinal cord (SC), motor axons that sort at the dorsal plexus (DP) to innervate the abductor (green) or adductor (orange) muscle are mixed within nerves 1-4. The C and D boxes denote the regions included in the maximum projections shown in C-D. The lateral view shows the arrangement of nerves 1-4 in the body wall (magenta) and innervation in the fin (green) as it is shown in C-D. The dashed line indicates where nerve 4 grows in the body wall behind the fin. B) Quantification of number of Schwann cells in the abductor or adductor innervation or associated with the plexus. n= 8 pectoral fins. C) Maximum projection of abductor innervation labeled with *Tubb:dsRed* and *37a> EGFP* to label Schwann cells. VP labels ventral plexus. Inset shows Schwann cell membranes that surround axon fascicles within the pectoral fin. Arrows point to axons that are not associated with Schwann cells. D) Maximum projection of body wall innervation of the same larvae shown in C. Nerves 3 and 4 are labeled. Inset shows dorsal plexus expanded in E-F. Asterisks label the dorsal region of abductor and adductor innervation. The rest of the fin innervation is not included in this maximum projection. E) Maximum projection of the dorsal plexus with a single plane shown in F. Abductor (Ab) and adductor (Ad) innervation and nerves 1/2 and 3 are labeled. The arrow in F points to Schwann cell membranes that completely wrap axonal fascicle in the plexus. Scale bars are 25 microns for C-D and 10 microns for the inset in C and E-F.

Following nerve injury, denervated Schwann cells also known as Bands of Bungner undergo morphological changes and can serve as substrates for regenerating axons^23,24^. We wondered whether a similar process occurs in the pectoral fin after axon injury. To address this question, we employed timelapse imaging during axon regeneration to observe Schwann cells labeled with *37A:Gal4, UAS:EGFP* and axons labeled with *Tubb:dsRed* **(Mov 3)**. Generally, the cell bodies of Schwann cells and their proximate membranes remained in place following axon injury, such that Schwann cell membranes reflect the pre-injury axon patterning footprint. Similar to what has been described in the trunk^23^, after axon fragmentation, elongated Schwann cell membranes changed from a tube-like morphology to a compressed morphology, likely due to the loss of axonal material. Moreover, distal membranes of Schwann cells became dynamic after axon denervation, repeatedly retracting and extending. Regenerating motor axons grew in close association with Schwann cell membranes, and the relationship between axons and Schwann cells is shown in **Fig 5 A-B (Mov 3)**, where Schwann cell membranes mark a choice point for regenerating axons. Here, a group of pioneer regenerating axons grew posteriorly along Schwann cell membranes in the dorsal musculature of the fin until they reached a choice point. Some of these axons continued to grow posteriorly, whereas others turned ventrally. Thus, within the pectoral fin, regenerating axons grow proximate to Schwann cells, consistent with the idea that the presence of Schwann cells might impact axon guidance at choice points.

**Figure 5:**
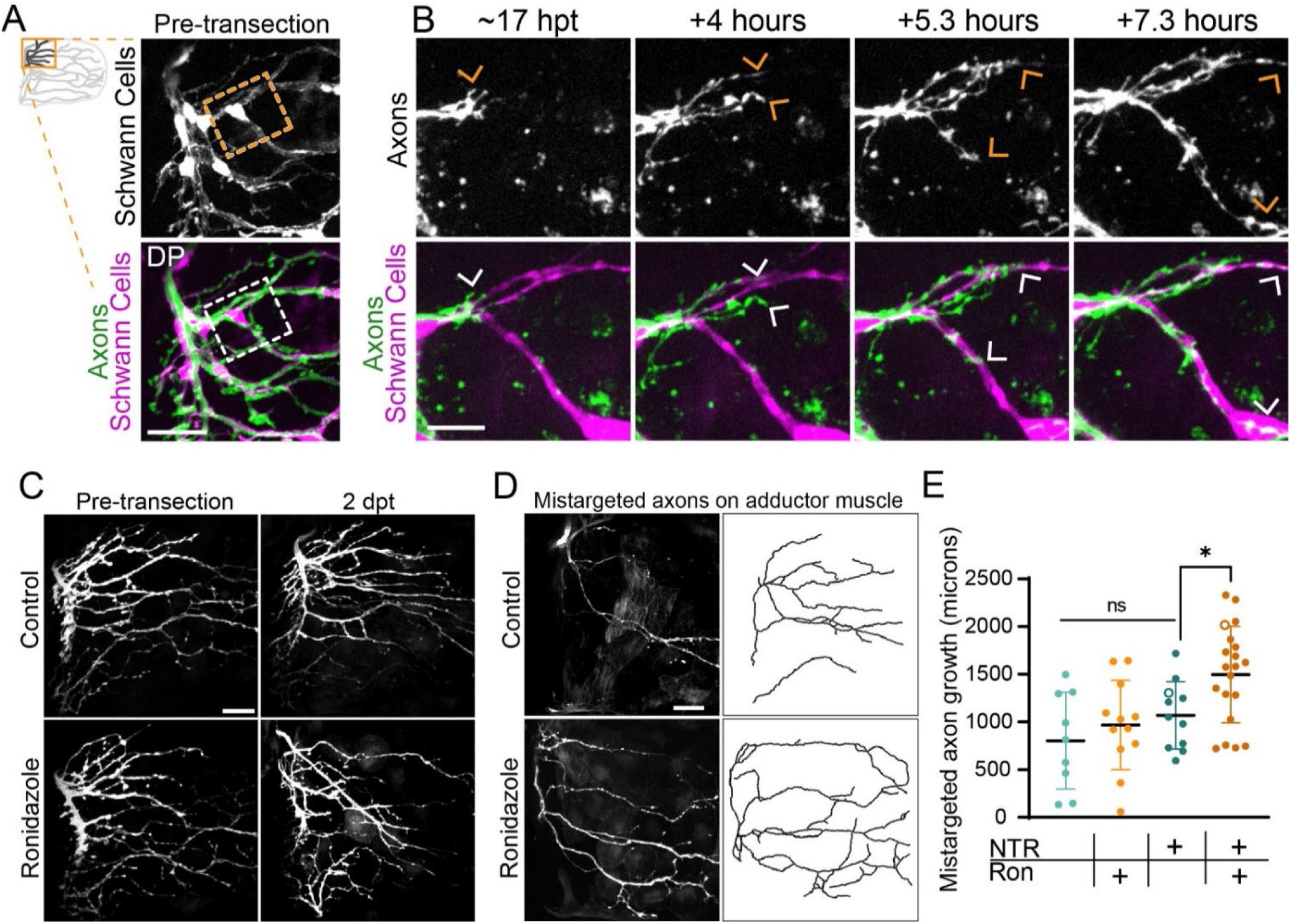
Schwann cells instruct axonal regrowth. A) Pre-transection maximum projection of axons labeled with *Tubb:dsRed* and Schwann cells labeled with *37a> EGFP*. Boxed inset outlines the region shown in B. DP labels dorsal plexus. B) Timelapse imaging captures pioneer axons (arrow) navigating a choice point. Axons remain tightly associated with Schwann cell membranes. C) Maximum projections of abductor-specific innervation labeled with *zCrest:GFP* in animals that also express *37a> NTR-tagRFP* before injury and at 2 days post transection (dpt). The addition of ronidazole ablated Schwann cells. D) Maximum projection and traces of mistargeted *zCrest:GFP-labeled* axons on the adductor muscle in *37a> NTR-tagRFP* animals treated with ronidazole or controls. E) Quantification of total amount of mistargeted zCrest:GFP+ axon growth on adductor muscle. Data points labeled with hollow circles denote the examples shown in B. *p<0.05, ns = not significant. one way ANOVA. Scale bars are 10 microns (B) and 25 microns (A, C-D).

### Glia promote axon targeting through a plexus

The high fidelity of axons targeting their original muscles (**Fig 2)** combined with the close association between regenerating axons and Schwann cells (**Fig 4)** led us to ask whether Schwann cells promote target-selective axon growth. To test this idea, we labeled motor neurons with *mnx1*:GFP to examine axon regeneration in *sox10(cls)^m241^* mutants^25^, which lack differentiated Schwann cells. We first analyzed the innervation patterning within the pectoral fin in *sox10* mutants compared to sibling controls prior to nerve transection. This revealed no difference between genotypes in the overall innervation pattern within the pectoral fin musculature **(Supp Fig 2A)**. In sibling controls, axons from nerves 1/2 and 3 were tightly fasciculated prior to and as they converged to form the “X” structure of the plexus (siblings: 3.4 ± 1.6 fascicles). In *sox10(cls)^m241^* mutants, prior to transection, axons at the dorsal plexus displayed a subtle increase in defasciculation (*sox10* mutants: 5.8 ± 1.75 fascicles; p=0.0025, t-test), suggesting that differentiated Schwann cells are required to promote or maintain axon fasciculation at the plexus during development **(Supp Fig 2B-C)**. Following transection of all fin motor nerves, regenerated axons in siblings re-formed the fasciculated “X” structure of the plexus, whereas *sox10* mutants displayed a marked increase in axon defasciculation (siblings: 8.2 ± 4 fascicles, *sox10* mutants: 15 ± 4.5 fascicles, p=0.0011, t-test) **(Supp Fig 2B-C)**. We qualitatively categorized the overall disorder of the plexus region as defined by axon defasciculation and axons entering the dorsal plexus region of the pectoral fin at ectopic locations. We found that compared to siblings, *sox10* mutant pectoral fins were more likely to exhibit a severely disorganized dorsal plexus (p=0.0001 Fisher’s exact test comparing categories Mild/Moderate with Severe in sibling controls (n = 13) and *sox10* mutants (n=11)) **(Supp Fig 2D)**. Thus, Schwann cells play a critical role at the dorsal plexus in promoting axon fasciculation.

To determine the role of Schwann cells in axon regeneration independent of development, we ablated Schwann cells specifically during axon regeneration. To do this, we chemogenetically ablated Schwann cells using the nitroreductase (NTR)/ronidazole system^26,27^.

Specifically, we used *37A:Gal4* to drive Schwann cell expression of *UAS:NTR-RFP*. First, we used the *37A:Gal4* driver to co-express *UAS:EGFP* and *UAS:NTR-RFP* and found that 95.7% of GFP-labeled Schwann cells expressed NTR-RFP (n = 95 GFP+ Schwann cells; 8 pectoral fins), demonstrating the feasibility of this approach to ablate most pectoral fin Schwann cells. We treated these larvae with ronidazole and then assessed Schwann cell ablation. Compared to untreated controls, within 12 hours of ronidazole treatment NTR-RFP-labeled Schwann cell membranes had retracted and cell bodies had rounded. After 24 hours, all NTR-RFP-labeled Schwann cells within the pectoral fin musculature were absent and RFP-positive debris was visible in the fin **(Supp Fig 3)**. We then assessed the role of Schwann cells specifically during axon regeneration by ablating Schwann cells after transecting fin motor nerves labeled by abductor-specific *zCrest:GFP*. We found that axons in Schwann cell-ablated pectoral fins regrew, demonstrating that Schwann cells are dispensable for axon growth towards and into the pectoral fin **(Fig 5C)**. However, while some *zCrest:GFP-labeled* axons were mistargeted to the adductor muscle in all fins (n = 52/52), there was an increase in the total length of mistargeted *zCrest:GFP-labeled* axon growth onto the adductor muscle in Schwann cell-ablated pectoral fins **(Fig 5 D-E).** Thus, Schwann cells are dispensable for axon regrowth but they are required during axon regeneration to promote target-selective axon growth in the pectoral fin.

Given the close association of Schwann cells in the plexus **(Fig 4 E-F)** and the increase in axon mistargeting between muscles after Schwann cell ablation, we asked how Schwann cells promote target-selective axon regeneration at the plexus. Prior to axon transection, *zCrest:GFP*-labeled axons formed a “Y” shape comprised of nerves 1/2 and 3 that converged at the dorsal plexus and then exclusively innervated the abductor muscle **(Fig 4E)**. In controls, at 2 dpt, regenerated *zCrest:GFP*-labeled axons formed a tightly fasciculated “X” structure, in which axons predominantly re-innervated the abductor muscle but a small fraction missorted at the plexus to innervate the adductor muscle **(Fig 6A)**. In contrast to controls, axons in Schwann cell-ablated pectoral fins failed to coalesce at a single plexus “X”. Instead, axons were defasciculated and entered both the abductor and adductor muscles at multiple entry points **(Fig 6B)**. We qualitatively categorized the overall disorder of the plexus region and found that compared to controls, Schwann cell-ablated pectoral fins were more likely to have a severely disorganized dorsal plexus (p=0.0003 Fisher’s exact test comparing categories Mild/Moderate with Severe in NTR-expressing animals treated with control (n = 11) or ronidazole (n=17)) **(Fig 6C)**. Thus, we demonstrate via both genetic and chemo-genetic cell ablation that during axon regeneration Schwann cells promote axon fasciculation to organize growth through the dorsal plexus, thereby preventing axon mistargeting onto the incorrect muscle.

**Figure 6:**
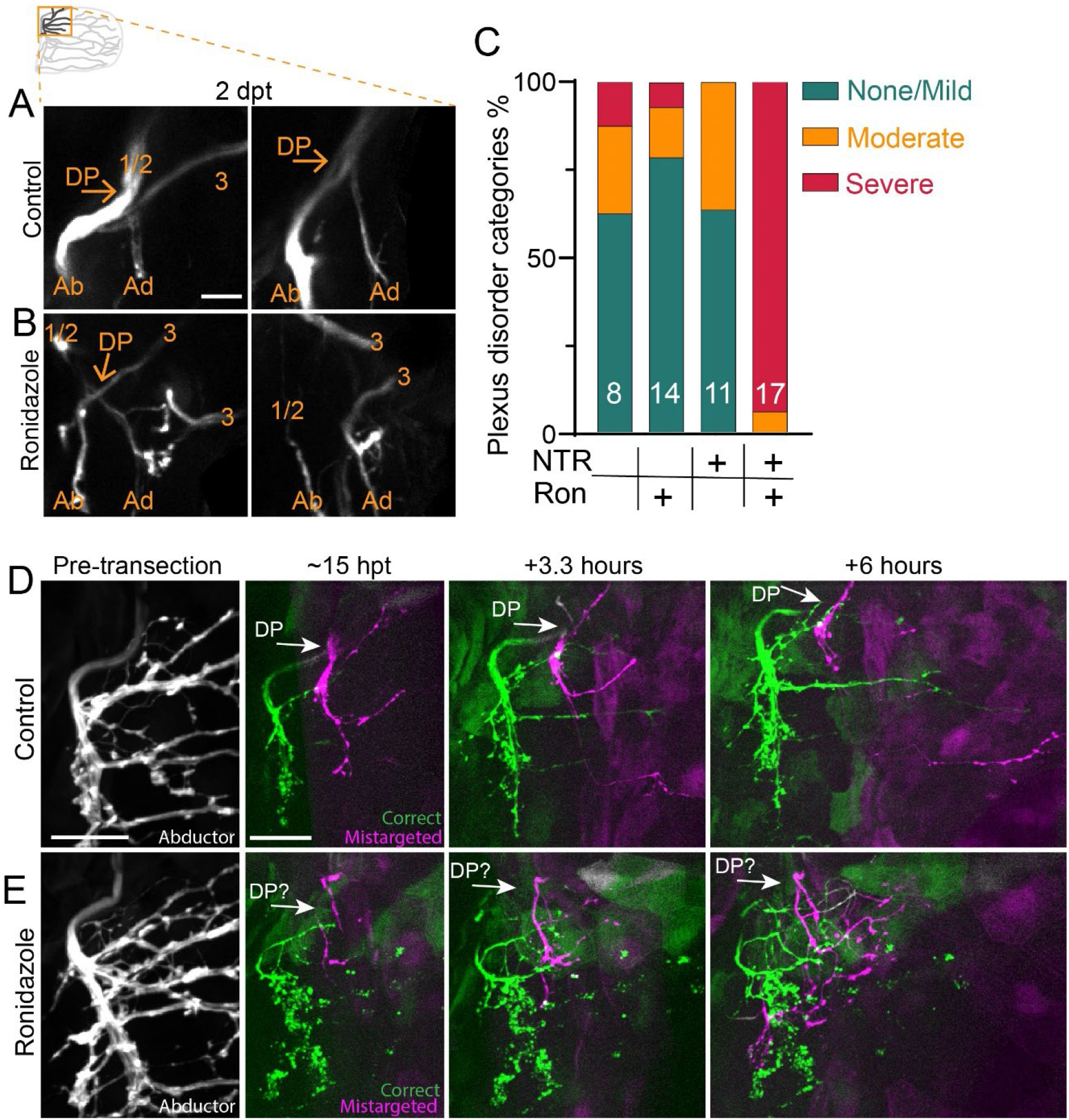
Schwann cells organize axon regeneration through the plexus. A) Maximum projection of the regenerated dorsal plexus at 2 dpt from two animals. Nerves 1/2 and 3, the dorsal plexus (DP), and the abductor (Ab) and adductor (Ad) innervation are labeled. Both were categorized as ‘none/mild’ plexus disorder. B) Two examples of plexus disorganization category ‘severe’ in animals with Schwann cells ablated after adding ronidazole. C) Categorization of plexus organization in regenerated fins. Numbers indicate n. NTR = nitroreductase, Ron = ronidazole. D-E) Timelapse imaging during regeneration of *zCrest:GFP* to label abductor-specific axons. Schwann cells express *37a> NTR-tagRFP* (not shown). D) In the control, the plexus is organized with fascicles correctly targeting the abductor muscle (green) and incorrectly targeting the adductor muscle (magenta). Axons extend orderly projections into the fin musculature. n=4/5 E) When Schwann cells are ablated after adding ronidazole, axon growth through the plexus is disorganized and axons are defasciculated. n= 4/4. Scale bars are 25 microns (D-E) and 10 microns (A).

Given the role of Schwann cells to organize axons at the plexus, we examined the dynamic nature of axon growth in this region. For this, we employed timelapse imaging of *zCrest:GFP*-labeled abductor-projecting motor axons in animals that also expressed *37A:Gal4; UAS-NTR* to ablate Schwann cells. By 15 hpt, axons in control fins have already sorted at the dorsal plexus and grew into the fin musculature (**Fig 6D)**. After exiting the plexus, axons that correctly targeted to the abductor muscle and axons that mistargeted to the adductor muscle remained tightly fasciculated prior to encountering the first branch points within the musculature (n=4/5). In contrast, in Schwann cell-ablated fins, axon growth was defasciculated and axons failed to converge at the plexus and instead grew aberrantly (n=4/4) **(Fig 6E, Mov 4)**. Combined, these results reveal that Schwann cells direct regenerating axons as they navigate towards and through the dorsal plexus.

## DISCUSSION

Regenerating axons re-establish connections to their original targets to achieve functional recovery. Target-selective axon regeneration is challenging to achieve when axons encounter stepwise choice points along their route. For example, after exiting the spinal cord, limb-innervating motor axons progressively converge and sort into target-selective bundles at the brachial plexus prior to innervating specific muscles. Here, we used the larval zebrafish pectoral fin to establish a genetically tractable model with unparalleled resolution to study target-selective axon regeneration through a plexus. We demonstrate that within two days following complete nerve transection, pectoral fin innervating motor axons navigate a series of stepwise choice points to regenerate with a surprising degree of high fidelity to their original muscle and muscle fiber targets. Through high-speed imaging of pectoral fin movements, we demonstrate that regenerated axons reform functional synapses. We use static and live imaging to show that axonal mistargeting occurs but that mistargeted axons are selectively retracted. In addition, we demonstrate that Schwann cells prevent axon defasciculation at the dorsal plexus and restrict axonal entry points into the pectoral fin, thereby promoting target-selective axon regeneration. Thus, our work reveals cellular mechanisms specific to regeneration that ensure appropriate axon targeting through a plexus.

### Fidelity of target-selection during regeneration

Functional recovery after peripheral nerve injury depends both on sustained axonal regrowth and on axons reinnervating their appropriate targets^28–31^. In rodent models, after complete nerve transection, regenerating motor axons preferentially reinnervate motor rather than cutaneous targets^32^. While these regenerating axons preferentially select their original nerve branch^33,34^, they frequently fail to accurately reinnervate their original muscle targets^35–38^. With a few exceptions^35^, determining the targeting precision of individual axons at the muscle or even single muscle fiber level has been challenging. In contrast to rodent models, in the larval zebrafish repeated and continuous imaging of labeled neuronal sub-populations or single neurons is accessible. Using this approach, we observed surprisingly high levels of fidelity by which regenerating motor axons re-innervated their original muscle (94.5%) and their original domain (85.5%). This specificity is consistent with previous reports on the specificity inferred through functional recovery in cichlid fish^39,40^, and on target fidelity in the larval zebrafish at the level of nerve branch and pharyngeal arch targets^8,41^. Thus, the zebrafish pectoral fin model combines a functional readout of recovery with single axon resolution, and also reveals that individual motor axons have retained significant capacity to reinnervate their original muscle fibers.

### Role of Schwann cells at the plexus

The dorsal plexus represents the first major choice point for pectoral fin motor axons, which converge at the plexus, intermingle, and sort between the abductor and adductor muscle. Compared to pre-transection sorting, we observed a marked increase in regenerating axons missorting at the dorsal plexus **(Fig 3)**. This suggests that regenerating axons make more sorting mistakes at the plexus, and/or that error correction is less effective in regenerating animals. This difference might reflect a change in the cellular or molecular composition of the plexus over time. Consistent with this notion, we uncovered a critical role for Schwann cells to prevent axon defasciculation and organize axon growth through the plexus specifically during regeneration **(Fig 5–6)**. Indeed, in both the zebrafish trunk and posterior lateral line, Schwann cells are dispensable for developmental axon outgrowth and targeting, but play important roles for axon regeneration^23,42^.

What are the mechanisms by which Schwann cells promote target-selective axon regeneration through the plexus? Given the concentration of Schwann cells at the dorsal plexus **(Fig 4)**, one possible mechanism is that Schwann cells play a structural role to effectively funnel regenerating axons to and through the dorsal plexus. In addition, it is likely that Schwann cells signal directly to regenerating axons to promote axon fasciculation and instruct axon growth through the plexus. Indeed, Schwann cells promote axon fasciculation via neuregulin signaling^43^ and are required in the larval zebrafish for fasciculation of regenerating posterior lateral line axons^42^. Moreover, there is precedence for Schwann cell-dependent regulation of axonal choice points, as a subset of Schwann cells upregulate the collagen *col4a5* to mediate axonal guidance decisions between dorsal vs ventral motor nerves in the larval zebrafish trunk via Slit/Robo signaling^8,9^. Which molecular cues might be expressed by Schwann cells at the dorsal plexus? Extensive work has characterized the role of several guidance cues, including polysialic acid and the cell adhesion molecule L1^44,45^, ephrinA-EphA4^46,47^, GDNF/Ret^48,49^, and Sema3A-Npn-1 signaling^50^ in developmental sorting of axons at a plexus. Yet whether these same cues promote axon sorting during regeneration, and whether they are expressed by Schwann cells, requires future investigation.

### Molecular logic of domain-specificity

After sorting at the plexus, fin innervating motor axons topographically target specific muscle domains. Here, we demonstrate that regenerating fin motor axons do not only reinnervate their correct musculature domains with fidelity, but they also return to their original muscle fibers **(Fig 2)**. Despite this specificity, the branching pattern of regenerated axons does not exactly match the developmental pattern, suggesting that axons may not exclusively reinnervate the exact same synapses **(Fig 2 A-B; Supp Fig 1)**. Pectoral fin motor axons form *en pessant* synapses across many muscle fibers and, likewise, individual muscle fibers are innervated by multiple axons^16,51^. Therefore, given the redundant innervation, it may not be necessary for axons to exactly reinnervate their original synaptic targets to regain coordinated fin movement.

The mechanisms that mediate topographic targeting of pectoral fin innervating motor neurons are unknown. In tetrapods, topographic motor targeting by lateral motor column neurons in the spinal cord is controlled by the distribution of EphA receptor expression in motor neurons and ephrin-A ligands in the limb^48^. In larval zebrafish, topographic targeting of cranial motor neurons to the pharyngeal arches, just anterior to the pectoral fin innervating motor neurons and the pectoral fin, respectively, is mediated by a retinoic acid gradient and Hgf/Met signaling during development^52^. Interestingly, Hgf/Met signaling is dispensable for targeting of these axons during regeneration, consistent with the idea that regeneration is not simply a recapitulation of development^41^. Which cells might mediate axonal target selection during regeneration? After injury, distal Schwann cells de-differentiate into repair Schwann cells, upregulate neurotrophic factors, and form bands of Bungner upon which regenerating axons grow (reviewed in^13,53^). Through live imaging, we observed regenerating axons growing adjacent to Schwann cells **(Fig 5B)**, which is consistent with Schwann cell processes guiding regenerating axons^54^. It is conceivable that within the pectoral fin, individual Schwann cells or other cell types could express a unique complement of guidance cues recognized by different neuronal populations to mediate local topographic targeting to fin domains. Future work, including identifying the molecular cues and relevant cell types that establish topographic targeting, will provide a comprehensive view of the mechanisms that mediate precise axonal targeting in the pectoral fin.

### Selective retraction of mistargeted axons

The nervous system uses several mechanisms to fine-tune axonal connections^55^. During development, such mechanisms include competition for growth factors that induces programmed cell death^56^, selective axon degeneration of missorted axons^57–59^, and selective axon retraction to refine axonal projections^60,61^. During regeneration, motor neuron collaterals of rat femoral nerves that incorrectly project to the skin are pruned, whereas projections to the muscle are maintained^34^, demonstrating that axons regrowing into the wrong tissue can be corrected. Yet, in comparison to development, the process of error-correction during axon regeneration, and whether it occurs within the target tissue, is less well-studied. Dynamic bouts of axon extension and retraction are common during axon pathfinding in development^62,63^ and in regeneration^8,9^. Similarly, in the pectoral fin, axons displayed probing behaviors during regeneration **(Mov 2-3)**. In addition to dynamic probing, we also observed that axons mistargeted to the incorrect muscle are selectively retracted while correctly targeted axons in the same fascicle remain **(Fig 3)**. This selective axon retraction of mistargeted axons occurred over large distances (>70 microns, at least half the width of the musculature) and so may be regulated by distinct mechanisms from dynamic growth cone probing. Ablation of Schwann cells specifically during axon regeneration revealed an increase in the length of mistargeted axons **(Fig 5 D-E)**, suggesting that Schwann cells may also play a role in axon retraction in addition to organizing axon growth at the plexus.

One important yet unresolved question is how are mistargeted axons recognized in the pectoral fin and what are the mechanisms that mediate their retraction? Through labeling the population of abductor-specific axons, we observed branches of labeled fascicles that had initially mistargeted onto the adductor muscle and later retracted, whereas other branches in the same mistargeted fascicle had stabilized and persisted. One intriguing possibility that could differentiate the mistargeted axons that retract vs those that persist is not whether they were on the incorrect muscle, but rather if they occupied the incorrect musculature domain. This would suggest that muscle-specific correction mechanisms would be concentrated at the plexus, where axons are actively sorting between muscles, while topographic domain-specific correction mechanisms would function within the fin musculature. Indeed, if being on the incorrect muscle was a cue for retraction, one would expect all muscle-mistargeted axons to be corrected. If, instead, there were domain-specific correction mechanisms in the fin, one would predict that an abductor-specific, dorsal domain targeting axon **(Fig 2E)** that mistargets to wrong muscle but still targets the ‘correct’ dorsal domain might be spared whereas a similar axon that mistargets to a different domain would be corrected. These possibilities can be distinguished through single axon labeling timecourse experiments. Through live imaging, we found that retractions of mistargeted axons initiated with strikingly consistent timing across pectoral fins (19 ± 2.6 hpt), perhaps suggesting that an unknown retraction factor is not expressed until this timing. Candidates for this retraction factor include nitric oxide and brain-derived neurotrophic factor, which mediate axon stabilization and retraction of retinal ganglia cells in the developing chick tectum^64^, and semaphorin signaling, which prunes branches of hippocampal neurons in the mouse^61^. Future studies are necessary to determine the molecular mechanisms that mediate the selective axon retraction we observe in regeneration pectoral fin axons. At a broader level, our data provide a powerful platform and a framework for future work to uncover the cellular and molecular mechanisms that mediate target-selective axon regeneration and error correction.

## ACKNOWLEDGMENTS

The authors thank members of the Granato lab for helpful feedback. We thank the UPenn Cell and Developmental Biology microscopy core and our fish facility staff. We are grateful to Shin-ichi Higashijima for communications regarding the zCrest:GFP transgenic line. This work was supported by the National Institutes of Health (K01NS119496 to L.J.W., NDRB from AMED to K. K., R01NS097914 and RO1 EY024861 to M.G.).

## AUTHOR CONTRIBUTIONS

L. J.W.: conceptualization, investigation, formal analysis, writing-original draft, visualization, funding acquisition

C.G.: investigation, formal analysis, writing-review & editing

K.K.: resources

M. G.: conceptualization, writing-review & editing, funding acquisition, supervision

## DECLARATION OF INTERESTS

The authors declare no competing interests

---

**Supp Fig 1:**
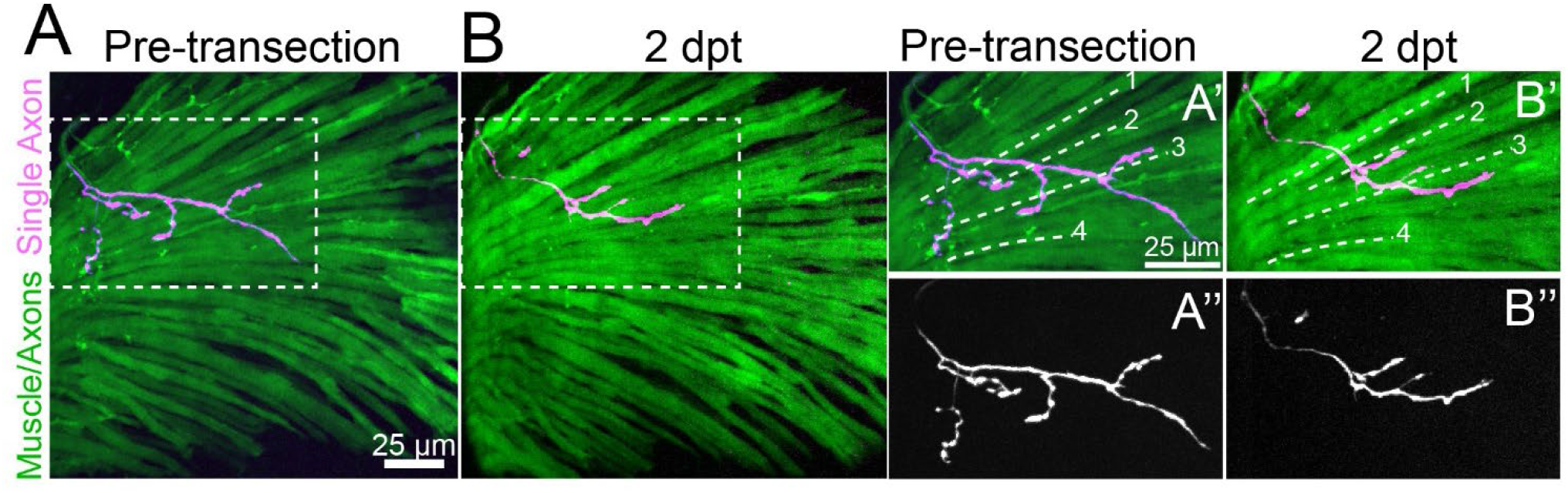
Motor axons regenerate to their original muscle fibers. A) Maximum projection prior to axon transection of abductor muscle in *Tg(α-actin:GFP);Tg(mnx1:GFP)* larvae showing muscles and motor axons (green motor axons are faint in this example) with a single axon trajectory in magenta. B) After 2 days, this labeled regenerated axon has reoccupied its original muscle fibers but has formed unique branches. Insets are expanded in A’ and B’ with individual muscle fibers labeled with the dotted lines to compare the axon location. The single labeled axon is shown in A’’ and B’’.

**Supp Fig 2:**
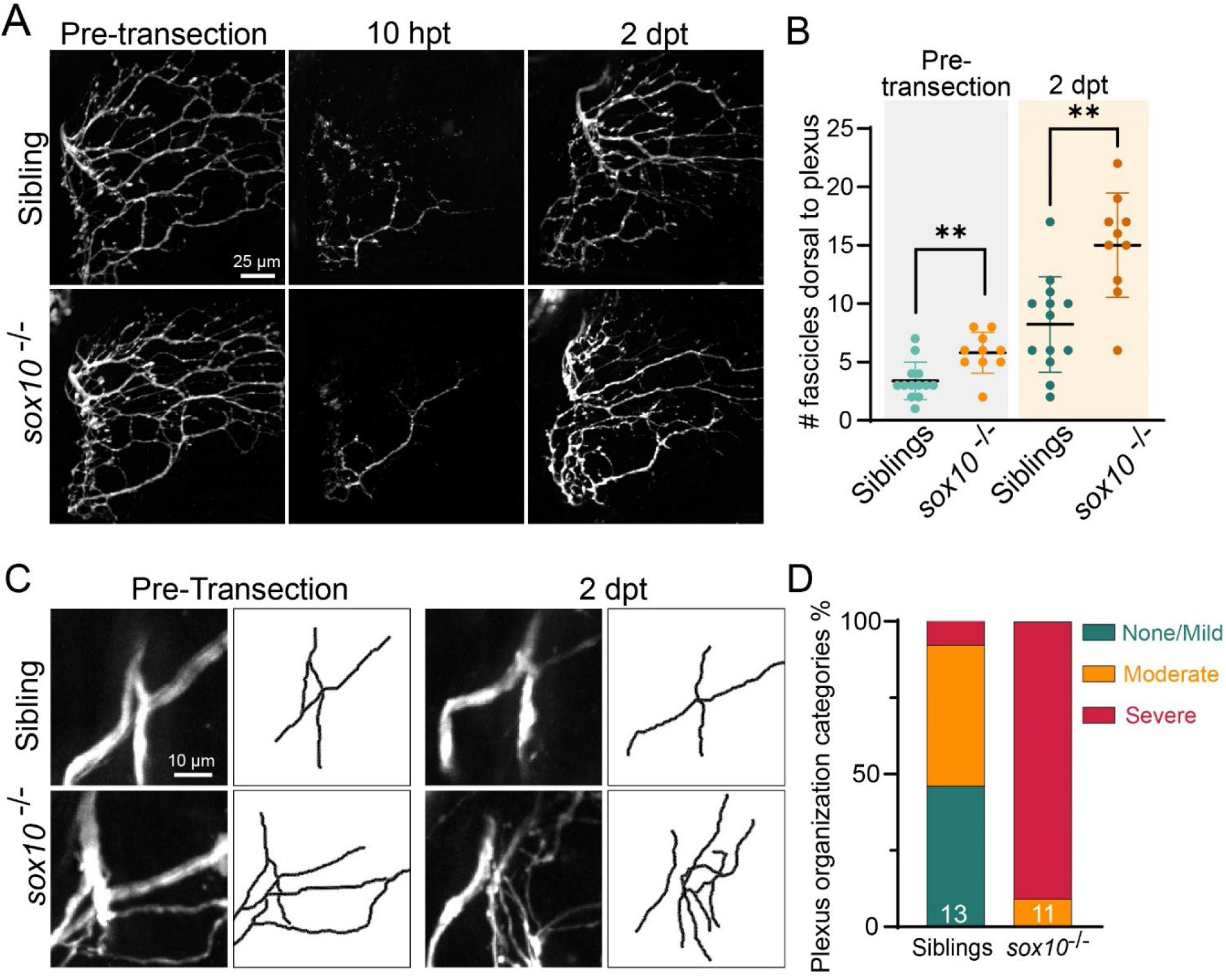
Differentiated Schwann cells organize axons at the plexus. A) Maximum projections of abductor muscle innervation labeled with *Tubb:dsRed* in sibling and *sox10^cls^* mutant larvae. The same fin is shown pre-transection, at 10 hpt (axon degeneration), and at 2 dpt (axon regeneration). B) *sox10* mutants display a subtle increase in axon fasciculation prior to axon transection that increases after axon regeneration C) Maximum projection through the plexus region before transection and at 2 dpt. *sox10* mutants display a disordered plexus region that worsens after injury. D) *sox10* mutants more frequently display severely disorganized axon patterning at the plexus at 2 dpt. ** p<0.01

**Supp Fig 3:**
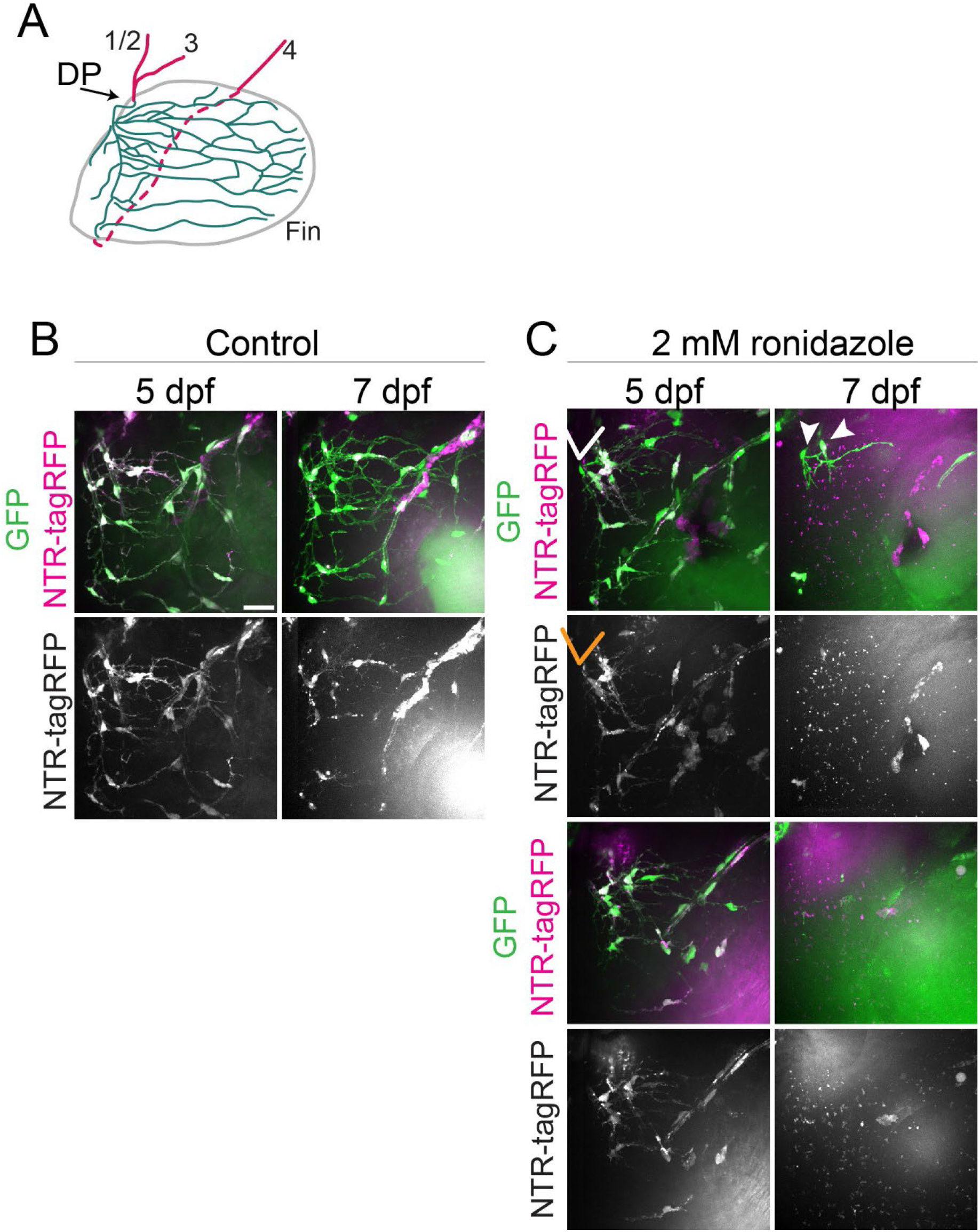
Ablation of Schwann cells. A) Schematic of innervation showing region included B-C. B) Maximum projection through pectoral fin of control *37a>EGFP, NTR-tagRFP* at 5 and 7 days post fertilization (dpf). C) Maximum projection through two examples of *37a>EGFP, NTR-tagRFP* larvae treated with ronidazole to ablate Schwann cells. The arrow points to a GFP-labeled Schwann cell that does not also express NTR-tagRFP and is spared from ablation at 7 dpf (arrowheads). Schwann-cell ablated animals display tagRFP-positive debris (puncta) at 7 dpf. Scale bars are 25 microns.

## MOVIE LEGENDS

**Movie 1:** Functional recovery of pectoral fin movements. High speed movies of pectoral fin movements were captured at 250 frames per second, but are slowed down 25x for this movie. The same fish was recorded for multiple timepoints. Prior to axon transection both pectoral fins move spontaneously. Nerves that innervate the right fin were transected while the left fin remained intact. At 5 hours and at 1 day post transection the right fin does not move. However, at 2 days post transection the right fin has regained movement.

**Movie 2:** Mistargeted axons are selectively retracted. *zCrest:GFP* labels axons that project only to the abductor muscle prior to axon injury whereas *Tubb:dsRed* labels all axons. The timelapse begins at 10 hpt with 15 minute intervals. In the first part, axon growth onto the adductor muscle is shown. Here, zCrest:GFP+ axons are incorrectly targeted to the adductor muscle and some of these axons are retracted while correctly-targeted Tubb:dsRed+ axons remain. In the second part, axon growth onto the abductor muscle of the same fin is shown. Correctly-targeted zCrest:GFP+ axons grow robustly.

**Movie 3:** Schwann cells instruct axonal regrowth. Pre-transection maximum projection of abductor innervation labeled with *Tubb:dsRed* and Schwann cells labeled with *37a> EGFP*. Timelapse movie begins ~18 hours after axon transection with 20 minute intervals between frames. The timelapse movie is first shown as a merged image but is followed by axon-only and Schwann cell-only movies. The orange arrows point to the example shown in figure 5B, with pioneer axons that navigate on or near Schwann cell membranes at a choice point. The arrow points to the first wave of pioneer axons which grow posteriorly. The filled orange arrowhead points to axons that turn at this choice point to grow ventrally. The white arrow points to an example of dynamic Schwann cell membranes.

**Movie 4**: Timelapse imaging during regeneration of *zCrest:GFP* to label abductor-specific axons. Schwann cells express *37a> NTR-tagRFP*. The timelapse begins ~15 hpt with 30 minute intervals between frames, showing a maximum projection of zCrest:GFP+ axons growing through and near the dorsal plexus during regeneration. In the control, axons are fasciculated to form the “X” shape of the dorsal plexus and extend orderly projections into the fin musculature. The orange arrow points to abductor axons while the pink arrow points to adductor axons. When Schwann cells are ablated after adding ronidazole, axon growth is disorganized and axons are defasciculated.

## STAR METHODS

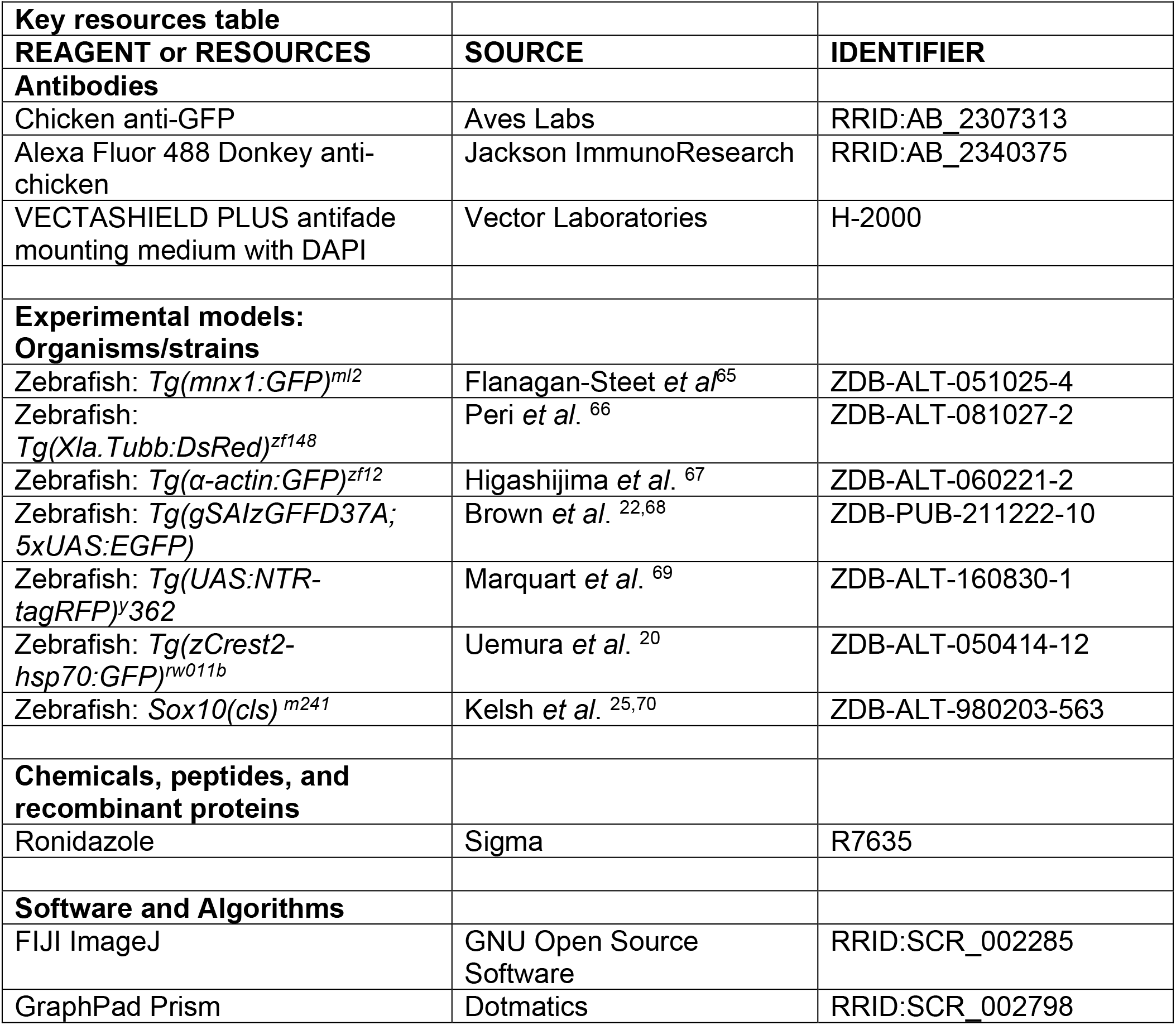

### Zebrafish strains and animal care

Protocols and procedures involving zebrafish (*Danio rerio*) are in compliance with the University of Pennsylvania Institutional Animal Care and Use Committee regulations. All transgenic lines were maintained in the Tübigen or Tupfel long fin genetic background and raised as previously described ^71^. The following transgenic lines and mutants were used: *Tg(mnx1:GFP)^ml2^* ^65^, *Tg(Xla.Tubb:DsRed)^zf148^* (referred to here as *Tubb:dsRed)* ^66^, *Tg(α-actin:GFP)* ^67^*, Tg(gSAIzGFFD37A; 5xUAS:EGFP)* (a kind gift from Dr. Sarah Kucenas) ^22,68^*, Tg(UAS:NTR-tagRFP)* ^69^*, Tg(zCrest2-hsp70:GFP)^rw011b^* (referred to here as *zCrest:GFP)*^20^, and *Sox10(cls)^m241^* ^25,70^. Homozygous *sox10(cls)* were identified by phenotype as mutants do not develop pigment. As our experiments in larval zebrafish occur prior to sex determination, sex was not a biological variable^72^.

### Motor axon transection

*mnx1:GFP, Tubb:dsRed*, or *zCrest:GFP* animals were immobilized with tricaine (MS-222) and mounted in 1.5% agarose with their right sides down on a glass-bottomed dish. The pectoral fin motor innervation was imaged prior to axon transection. Motor axons were transected using an Ablate! 532 nm attenuable pulse laser (Intelligent Imaging Innovations (3I), Denver, CO) in the location shown in Fig 1C, with care taken to transect axons at least 15 microns away from the dorsal plexus so the plexus was not damaged and at a low laser power to not cause substantial damage to the surrounding tissue. Animals with significant tissue damage, evident by immediate fragmentation of axons in the region and rippling or movement of the surrounding tissue, were excluded. All motor nerves innervating the fin were transected. Nerves were considered transected when a disruption in the fluorescent signal was obvious at the transection site and when blebbing was present within the distal, transected axons in the fin (within 30 min but fragmentation is obvious by 3-5 hours). Animals were unmounted and housed in single wells of a 12- or 24-well dish at 28 degrees and re-mounted for imaging or behavioral testing as indicated.

### Sparse neuronal labeling

A DNA vector encoding *mnx1:mKate* was injected as previously described^73,74^ into one-cell-stage *mnx1:GFP* or *mnx1:GFP; α-actin:GFP* embryos. Embryos were screened at 1-3 dpf for larvae expressing mKate sparsely in the anterior spinal cord. At 5dpf, nerves were transected as described above if they had sparse mKate-expressing axons innervating the pectoral fin. Images were taken from 5-8 hpt to confirm fragmentation. Sparsely labeled axons were scored blind before injury and after regeneration for the muscle and domain the distal end of the axons occupied. Labeled axons were included in the analysis if they had a distinct localization that was confined to a domain of the fin that was distinguishable from other axons (for example, a single fin might have labeled axons that innervate domain 1 on the abductor muscle and domain 4 on the adductor muscle; these would be scored separately).

### Time-lapse imaging

Larvae were anesthetized with tricaine and mounted in 1.5% agarose on a glass bottom dish. For live imaging of Schwann-cell ablated animals, larvae were mounted in a chamber slide allowing for controls to remain in E3 media while ablated animals were maintained in E3 media plus 2 μM ronidazole. Animals were timelapsed using a 40x or 60x lens on an ix81 Olympus spinning disk confocal in a temperature chamber set to 28°C as previously described^75^. Stacks through the pectoral fin were captured in 1.5 μm slices with 15–30-minute intervals. Animals were imaged continuously for up to 3 days.

### Schwann cell ablations

*Tg(gSAIzGFFD37A; UAS:NTR-RFP; zCrestHSP70:GFP)* ^22,68^ larvae were anesthetized with tricaine, pectoral fins were imaged, and motor axons were transected. Larvae were unmounted into 12 well dishes and placed in E3 with 2uM ronidazole at 28 degrees. Control animals remained in E3 media. The ronidazole was replaced every day until the end of the experiment. Larvae were re-imaged at 1 and 2 days post transection. To assess the efficacy of Schwann cell ablations, *Tg(gSAIzGFFD37A; UAS:NTR-RFP; UAS:EGFP)* animals were imaged and treated with ronidazole or control media in parallel with transection experiments.

### Whole-mount immunohistochemistry and imaging

To label Schwann cell nuclei, after treatment with ronidazole for 2 days *Tg(gSAIzGFFD37A; UAS:NTR-RFP; UAS:EGFP) or Tg(gSAIzGFFD37A; UAS:NTR-RFP; zCrestHSP70:GFP)* animals were fixed for 1 hour at room temperature in sweet fix (4% paraformaldehyde with 125mM sucrose in PBS) plus 0.1% Triton X-100 (Fisher, BP151). Animals were washed in phosphate buffer and incubated overnight at 4°C in primary chicken anti-GFP antibody (1:2000, Aves labs, GFP-1010) in incubation buffer (2 mg/mL BSA, 0.5% Triton X-100, 1% NGS). After washing in phosphate buffer, animals were incubated overnight at 4°C in Alexa Fluor 488 donkey anti-chicken secondary antibody (1:1000, Jackson ImmunoResearch, 703-545-155) in incubation buffer. After washing with phosphate buffer saline, larvae were incubated overnight at 4°C in Vectashield plus with DAPI (Vector Laboratories, H2000). Animals were mounted in agarose in a glass-bottomed dish and imaged in 1.5 μm slices using a x40 water immersion lens on a Zeiss LSM880 confocal microscope using Zen software. These samples were used to count total Schwann cell numbers.

### Functional recovery

Larvae were tested for fin movements and then were immediately mounted for imaging and/or nerve transection. The same larvae were tested repeatedly throughout the experiment. Larvae were acclimated to room temperature for 15 minutes prior to behavior testing. Spontaneous fin movements were recorded at 250 frames per second using a high-speed camera mounted on a stereomicroscope and Photron Fastcam Viewer software. Spontaneous movements were defined as alternating, rhythmic pectoral fin movements in which the fin extends maximally perpendicular to the body wall. Bouts in which the larvae was moving, turning, or startled, in which pectoral movements can extend almost parallel to the body wall, were excluded. Maximum fin displacement angles were measured in Fiji during one bout of spontaneous fin movement per fish per timepoint. Data were excluded for some of the 5 hr and 24 hr timepoints if the location of the non-moving fin could not be distinguished from the bodywall of the larvae.

### Imaging processing and quantification

Fluorescent signal from the abductor vs adductor innervation was manually separated using Fiji^76^ as previously described^51^. Briefly, channels were separated, the background was subtracted, channels were merged, and the image was converted to RGB or 8-bit tiff. Using the 3D viewer, stacks were rotated to a dorsal view so that the innervation from the abductor vs adductor muscle was distinct. Axon signal was removed by selecting and filling a region, resulting in the corresponding area being filled with the background color. This process was completed separately for the abductor or adductor innervation and the corresponding signal was merged back together if necessary. Figures show abductor or adductor innervation or a merge of the two as noted. For most figures, brightness and contrast were adjusted individually for each image, so intensity of signal should not be compared. All quantification was blind to genotype or treatment. To quantify mistargeted axonal growth **(Fig 5D-E)**, the adductor innervation was isolated and zCrest:GFP-positive axons were traced using Simple Neurite Tracer^77^ in Fiji. To quantify plexus organization, 3D stacks were categorized based on the structure of the dorsal plexus as labeled by zCrest:GFP. Plexuses were categorized in the following way: ‘None/Mild’ category had tightly fasciculated axons that cross to form an “X” shape, with a thick abductor fascicle and a thinner adductor fascicle, ‘Moderate’ category had some defasciculation but still formed the “X” shape, ‘Severe’ category had severe defasciculation, extraneous fascicles that enter the ‘plexus’ region via ectopic routes, and/or failed to form the “X” shape.

### Statistical analysis

Data were imported into GraphPad Prism for analysis. Statistical analysis was performed as indicated throughout the text.

## Notes

### Competing Interest Statement

The authors have declared no competing interest.

